# The oldest freshwater crabs: claws on dinosaur bones

**DOI:** 10.1101/747535

**Authors:** Ninon Robin, Barry W.M. van Bakel, Matúš Hyžný, Aude Cincotta, Géraldine Garcia, Sylvain Charbonnier, Pascal Godefroit, Xavier Valentin

## Abstract

With approximately 1,500 extant species, freshwater crabs (Decapoda: Brachyura) are among the most diverse decapod crustaceans. Nevertheless, their fossil record is extremely limited: only Potamidae, Potamonautidae and Trichodactylidae are reported up to the Eocene of the Neotropics so far. This work documents unusually large decapod claws from the Upper Cretaceous (Campanian) continental deposits of Velaux and vicinity (southern France), in close association with large vertebrate remains. In addition to (1) the systematic assignment of these claws, the study addresses (2) the salinity trends in the deposit environment from its faunal assemblage and the elementary chemical patterns of fossils, and (3) the likely scenario for their auto/allochtony in the Velaux fluvial system. These claws belong to a new taxon, *Dinocarcinus velauciensis* n. gen. n. sp., referred to as Portunoidea sensu lato, a group of “true” crabs nowadays linked to marine systems. However, the faunal assemblage, the claw taphonomy and the carbonates Y/Ho signatures support their ancient freshwater/terrestrial ecology, making them the oldest reported continental brachyurans and extending the presence of crabs in freshwater environments by 40 Ma. Either as primary or as secondary freshwater crabs, the occurrence of these portunoids in Velaux is an evidence for the independent colonizations of continental environments by multiple brachyuran clades over time, as early as the Campanian.

## I. Introduction

With approximately 1,500 extant species^1^, freshwater brachyuran crabs (Decapoda: Brachyura) are among the most diverse decapod crustaceans. Nevertheless, their fossil record is extremely limited. Representatives of three families were identified unequivocally as fossils, including Potamidae Ortmann, 1896^2^, Potamonautidae Bott, 1970^3^ and Trichodactylidae H. Milne Edwards, 1853^4 5–7^. Articulated exoskeletons of fossil freshwater crabs are rare^7–14^, although isolated cheliped fingers are much more frequent, but difficult to evaluate taxonomically^9,15–18^. Up to now, *Tanzanonautes tuerkayi* Feldmann et al., 2007^13^ (Potamonautidae) from the Oligocene of Tanzania (*ca* 30 Ma) is the oldest fossil record of a freshwater brachyuran in the Old World and there is no remain of Potamidae older than early Miocene^9^. The oldest record of freshwater crabs is from the middle Eocene of the Amazon Basin (*ca* 40 Ma) and belongs to the family Trichodactylidae^7^, a group of crabs that likely colonized freshwater habitats independently from potamoids, as indicated by morphology^19^ and molecular phylogeny^20^.

The present paper reports the remains of brachyuran crabs from fluvial Late Cretaceous (late Campanian; *ca* 72-74 Ma) localities of southern France (Velaux-La Bastide Neuve and vicinity), fossilized in associations with vertebrate remains. Close associations of different and diverse fossil organisms may both (1) be the result of a long-distance transport of allochthonous remains or (2) testify of local biocoenoses for which members of quite restricted ecosystems deposited altogether. These claws are of exceptional large size compared to most Late Cretaceous marine crab claws; and interestingly do not conform the morphology of any extant freshwater crab family. The presence of presumably freshwater crabs in Campanian deposits is quite unexpected and represents the earliest record of the colonization of freshwater environments by brachyuran decapod crustaceans. It roughly doubles the previously oldest evidence of 40 Ma, and would further support the independent invasion of freshwater environment by several distinct brachyuran lineages^6,21,22^.

The herein study aims at (1) proposing an accurate systematics assignment for these claws, (2) characterizing the actual salinity trend of their deposit environment, based on the channel fauna assemblage and elementary chemical patterns of fossils and (3) identifying the relevant taphonomic scenario for the presence of crab claws within a fluvial system. As all these approaches support a neat freshwater or terrestrial signature for the living paleoenvironment of these large-clawed brachyurans, we then discuss the implications for presumed multiple invasions of freshwater habitats by crustacean decapods over time.

## II. Velaux-La Bastide Neuve channel

Velaux-La Bastide Neuve is located in the western part of the Aix-en-Provence Basin, Southeastern France. K-Ar dating of the locality was attempted based on glauconites collected from sandstones^23^, but these minerals are clearly reworked from lower Aptian marine limestones and are therefore useless for dating the site^23^. Magnetostratigraphic analysis of the deposits, however, correlates with the normal chron of chron 32^23^, corresponding to an age of 71.6 to 74 Ma^24^. Along with correlations with charophytes and dinosaur eggshell biozones, a late Campanian age for the locality may confidently be proposed^23,25–27^. The fossil site is mostly known for its vertebrate assemblage, recovered from three different sedimentological sequences and corresponding to newly described dinosaur (titanosaurid sauropod^26,28^; rhabdodontid ornithopod^29^) and pterosaur (azhdarchid pterosaur^30^) taxa, as well as eusuchian crocodilians^27^. Apart from the diapsids, vertebrates consist of disarticulated pleurodiran and cryptodiran turtles, disconnected remains of sarcopterygian and actinopterygian fishes and, chondrichthyan teeth. Freshwater bivalves (*Unio*) and gastropods (*Physa*, *Melania*)^30^, macro-remains of angiosperm plants and charophytes complete the whole fossiliferous assemblage together with the herein described crustacean remains. The lithological section consists of 16.3 meters of alternating sandstones, siltstones – including paleosols – and mudstones. Lacustrine limestones occur in the uppermost part of the section. The succession was deposited in a fluvio-lacustrine environmental setting. The sedimentology of the site together with the fossil assemblage indicates a likely freshwater setting for the deposits. The succession of conglomeratic sandstones, siltstones (including paleosols), mudstones and lacustrine limestone on top of the stratigraphic section indicate sedimentation in, respectively, a low-energy fluvial channel, channel levees, alluvial plain and lake^23^. Given the proximity of Velaux to the paleo-coast during the Late Cretaceous^31,32^, occasional marine incursions are not excluded, even though they were not recorded at Velaux-La Bastide Neuve nor at other fossiliferous localities of the same age in the region^23,25^.

## III. Results

The studied material (Tab. 1) consists of seven (partial) claws and associated vertebrate remains collected from sequence 2 of the sedimentary succession of Velaux-La Bastide Neuve locality and one from the close locality of Rognac-Les Frégates (about four km from Velaux, corresponding to similar layers). Specimens are housed in the palaeontological collections of the municipal paleontological and archeological structures of Velaux (Musée du Moulin Seigneurial/Velaux-La Bastide Neuve: MMS/VBN.00.004, 09.69e, 12.A.006, 02.94, 09.43, 09.132d, 12.A.003) and of the Muséum d’Histoire naturelle d’Aix-en-Provence, France (MHN AIX PI 1991.1, coll. Valentin).

**Table I.** Examined fossil material and preservation state.

| Catalogue | Number | Item | Preservation | Type status |
| --- | --- | --- | --- | --- |
| MMS/VBN. | 00.004 | left-hand claw, with dactylus | well preserved | Holotype |
| MMS/VBN. | 09.69e | right-hand fixed finger | moderately preserved | / |
| MMS/VBN. | 12.A.006 | right-hand fixed finger | moderately preserved | Paratype 4 |
| MMS/VBN. | 02.94 | right-hand fixed finger | well preserved. very large | Paratype 1 |
| MMS/VBN. | 09.43 | right-hand fixed finger | coarse, surface partly dissolved | / |
| MMS/VBN. | 09.132d | right-hand fixed finger | well preserved | Paratype 2 |
| MMS/VBN. | 12.A.003 | right-hand dactylus | moderately preserved | Paratype 3 |
| MHN AIX | PI 1999.1 | left-hand claw | well- preserved | / |

### III.I. Systematic paleontology

#### III.1a: The crabs

##### Preliminary remarks

Taxonomic assignment of isolated claws of brachyuran crabs at the species or genus level is difficult, if not impossible in many cases. If direct comparisons with extant taxa is straightforward and often helpful for identifying Pliocene and Pleistocene brachyurans^33–36^, this tasks already becomes more complicated when working with material of Miocene age^37–40^. The taxonomic evaluation of isolated fossil cheliped fingers is of course further challenging. Erecting new taxa based on isolated brachyuran chelae alone has not been attempted yet, although this method is regarded as valid in other decapod groups, including paguroid hermit crabs^41–43^, erymid lobsters^44,45^ or callianassid ghost shrimps^46,47^. If working with distinct claw morphologies, the erection of new taxa can be done^48^. Alternatively, parataxonomy can be used^49^. We assume that the morphology and also the size of the studied claws from the two localities Velaux-La Bastide Neuve and Rognac-Les Frégates are distinct enough to warrant the validity of the new form genus *Dinocarcinus*. Based on the general morphology of its claws, we include this form genus within Portunoidea sensu lato^50^. Taxonomic characters in the *Dinocarcinus* material do not allow their classification within any known family, but rather point out its affinities to portunoids. In this sense, *Dinocarcinus velauciensis* is kept in open nomenclature.

Portunoidea sensu lato (see above)

*Dinocarcinus* n. gen. Van Bakel, Hyžný, Valentin & Robin Figs. 1, 2, 3

**Figure 1.**
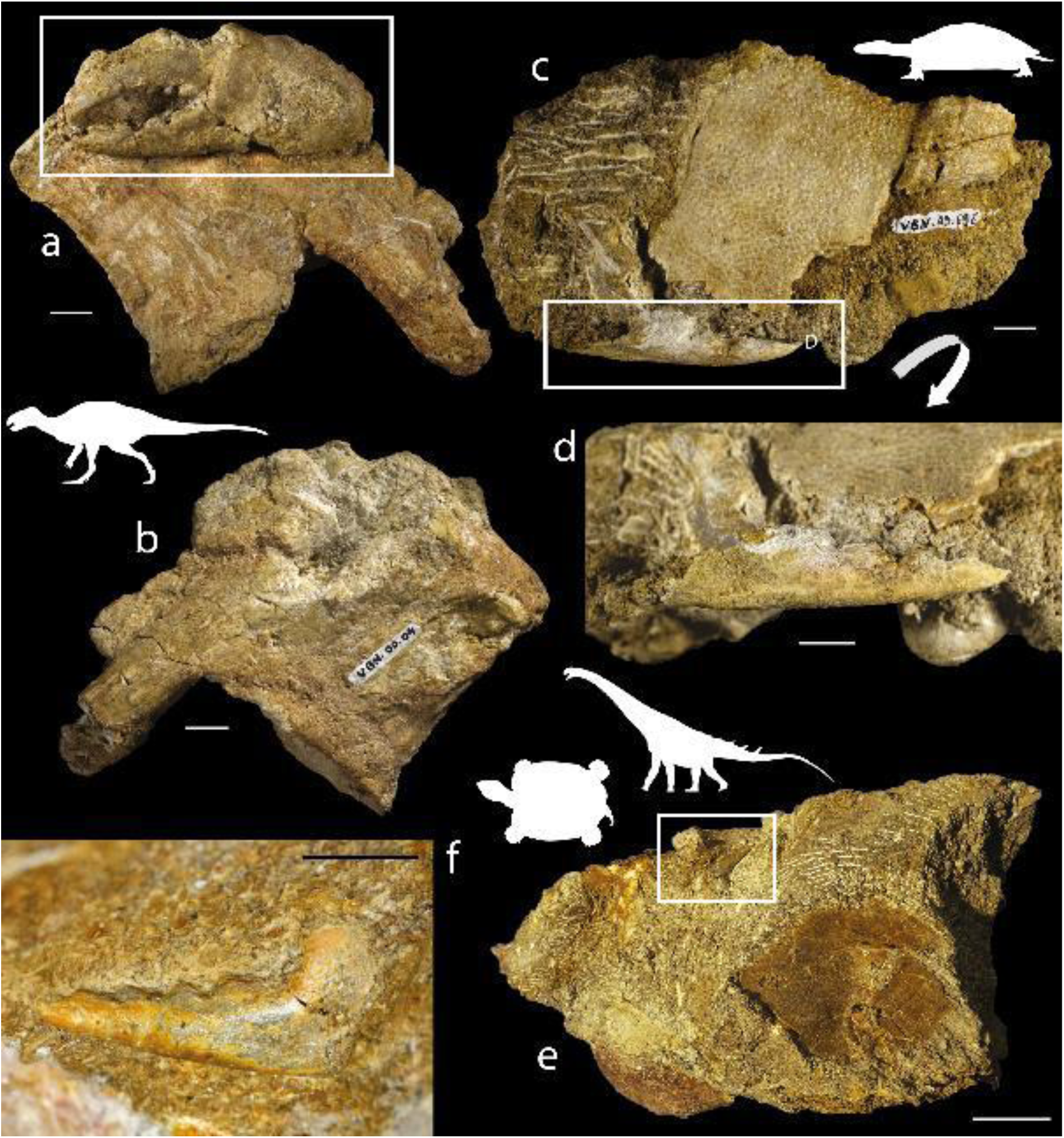
*Dinocarcinus velauciensis* Van Bakel, Hyžný & Robin n. gen., n. sp. Claws associated with vertebrate remains. **a-b**. MMS/VBN.00.004: anterior (a) and posterior (b) views of the claw-bearing ornithopod (rhabdodontid) vertebra, **c-d**. MMS/VBN.09.69e: outer view (c) and close-up (d) of the close sedimentary association of the claw with a turtle (solemydid) plastral plate, **e-f**. MMS/VBN.12.A.006: outer view (c) and close-up (d) of the block association of the claw with a turtle (bothremydid) plastral plate, ornithopod (rhabdodontid tooth) and partial sauropod (titanosaurid) dorsal vertebra with its ossified tendon. Scale bars = 1 cm (a-d, f); = 3 cm (**e**). Photographs. L. Cazes.

##### Etymology

Denoting the actual association with dinosaur (ornithopodan) remains.

##### Type species

*Dinocarcinus velauciensis* n. gen. n. sp.

##### Diagnosis

Chelae large and massive. Fingers gaping, arched, with strong teeth, proximal tooth molariform. Fixed finger dorsal surface with single “pitted groove”, palm surface smooth, articulation with dactylus oblique, prominent.

##### Remarks

The morphology of the claws, namely heavily calcified fingers, strong molariform teeth and grooved fingers, are typical for some eubrachyuran crabs. There are only few representatives of podotreme clades which grew to big sizes, with particular exceptions of Dakoticancridae and Cenomanocarcinidae, which could have claws comparably large to *Dinocarcinus velauciensis* n. gen. n. sp. Figured claws of *Avitelmessus* Rathbun, 1923^51^ show the following^51–53^: the fingers’ (distal) teeth are not molariform, fingers are less robust and less strongly calcified than the remains studied herein. The claws of *Avitelmessus* are more curved, the palm is longer than the fingers, both palm and fingers have crests and grooves; the fingers tips are hooked; distinguishing it easily from *Dinocarcinus* n. gen. The palaeocorystoid Cenomanocarcinidae could attain large sizes and had massive claws. The claws of *Cenomanocarcinus* Van Straelen, 1936^54^ are figured by Guinot et al.^55^ (fig. 6) and are characterized by spinose claws, flattened in cross section, a slightly downturned fixed finger, spines along the upper margin of the claw and dactylus, and hooked tips. Compare also the very large claws of ‘*Oncopareia’ heterodon* Bosquet, 1854^56^, now considered to be a palaeocorystoid (in Jagt et al.^57^: plate 5). As discussed above, the claw morphology of *Dinocarcinus velauciensis* n. gen. n. sp. does not match that of the Dakoticancroidea, Palaeocorystoidea, or any known Podotremata.

Within Eubrachyura, the robust, strongly calcified fingers, overall claw shape, and molariform teeth, match that of the Portunoidea. This large group of overall large-sized crabs have several Mesozoic occurrences, and some of them in large sizes. *Ophthalmoplax* Rathbun, 1935^58^, now considered a representative of Macropipidae^59^ has a great size range, from very large *Ophthalmoplax brasiliana* Maury, 1930^60^ to rather small *O. minimus* Osso et al., 2010^61^. Their claws [compare^62^ (figs. 3.2, 3.3, 4.2, 4.13, 4.14) with^61^ (fig 6.7)] are spinose, keeled, with major claws showing a large bulbous proximal tooth ascribed to shell breaking mechanisms^63^. These specialized claws can be easily distinguished from the more simple, unarmed claws of *Dinocarcinus* n. gen. *Eogeryon* Osso, 2016^64^ (Cenomanian of Spain) is assigned to the Portunoidea in its own family (Eogeryonidae Osso, 2016^64^). Geryonidae Colosi, 1923^65^ and Eogeryonidae are considered as early diverging families within Portunoidea. *Eogeryon* is characterized by large claw size relative to the carapace, equal ratio palm-fingers, with strongly calcified fingers with molariform teeth, and grooved fixed finger. Its claw morphology is typical of that of Portunoidea, and compared with that of *Styracocarcinus meridionalis* (Secrétan, 1961^66^) from the ?Campanian, of Morocco. Claws of *Litoricola macrodactylus* (Van Straelen, 1924^67^) from the Paleocene of southern France and Northern Spain, are highly comparable with those of *Dinocarcinus* n. gen., however they show a bulbous proximal crushing tooth on the dactylus of the major claw. Also, the fingers in *Dinocarcinus* n. gen. are more gaping as in *Litoricola*.

The claw morphology of *Dinocarcinus* n. gen. shows few diagnostic characters for superfamily level assignment (Portunoidea), namely heavily calcified claws, a grooved fixed finger, molariform teeth, palm and fingers subequal in length, and blunt, non-hooked fingertips (Fig. 2). More accurate assignation is not possible at this point. An early diverging position within Portunoidea is possible considering morphology, geologic age, large size, and similar families occurring at that time.

**Figure 2.**
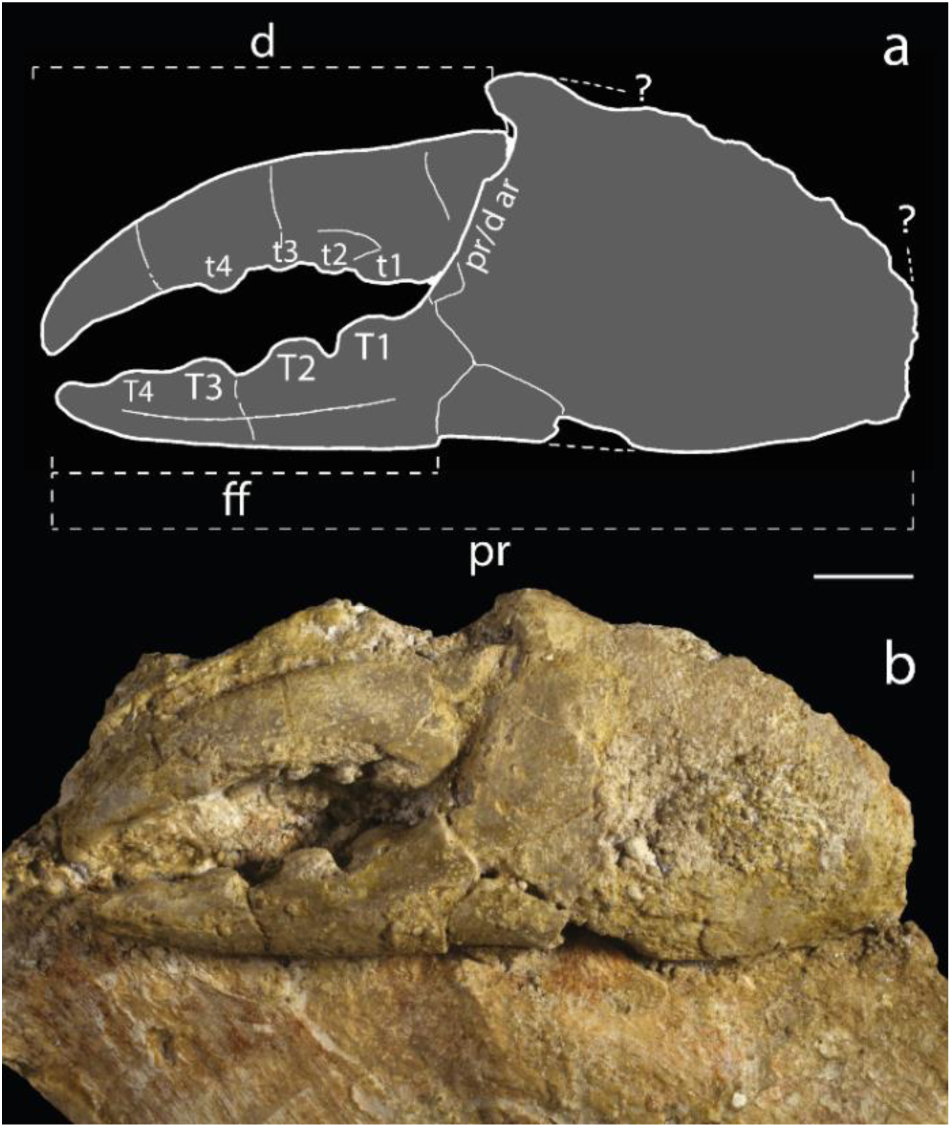
*Dinocarcinus velauciensis* Van Bakel, Hyžný & Robin n. gen., n. sp. **a-b** Illustrative features of holotype MMS/VBN.00.004 (complete chela). d = dactylus; ff = fixed finger; pr = propodus; pr/d ar = propodus/dactylus articulation; t1-4 = dactylus teeth; T1-4 = fixed finger teeth; ? = questionable limits. Scale bars= 1 cm. Drawing. B. van Bakel, photograph. L. Cazes

*Dinocarcinus velauciensis* Van Bakel, Hyžný, Valentin & Robin n. sp. Figs. 1, 2, 3

**Figure 3.**
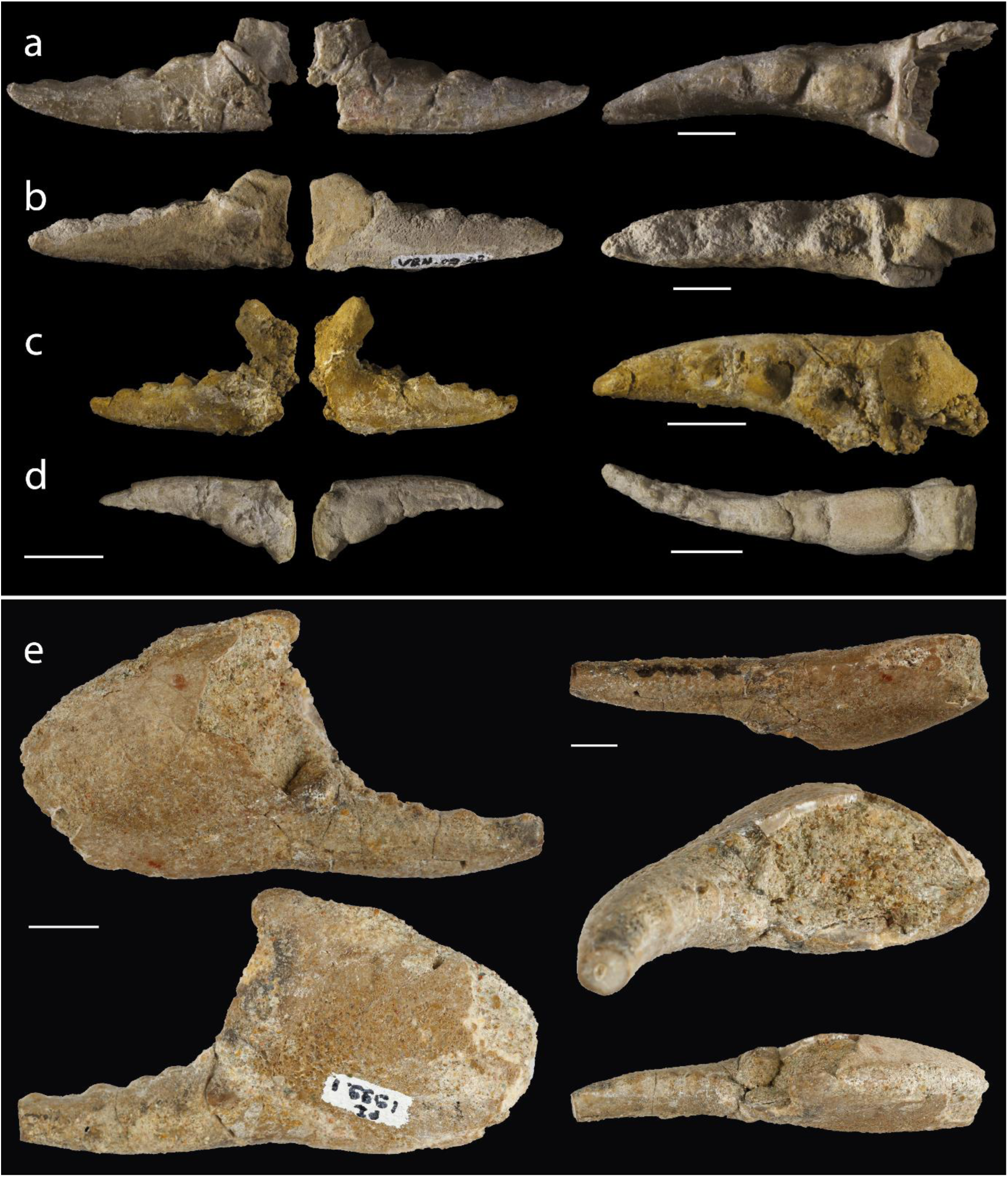
*Dinocarcinus velauciensis* Van Bakel, Hyžný & Robin n. gen., n. sp. Isolated claws found scattered in sediment**. a-d.** From left to right: inner, outer and occlusal views of dactyli showing teeth arrangement. **a.** MMS/VBN.02.94, **b.** MMS/VBN.09.43, **c.** MMS/VBN.09.132d **d.** MMS/VBN.12.A.003. **e.** MHN AIX PI 1991.1 (coll. X. Valentin) from the Campanian of Rognac, 4 km from Velaux. From up to down and left to right: inner, outer, marginal and occlusal two views showing teeth arrangement. Scale bars = 1 cm. Photographs. L. Cazes (a-d), Y. Dutour and E. Turini (e).

##### Type material

Holotype: MMS/VBN.00.004; Paratypes 1-4: MMS/VBN.02.94, 09.132d, 12.A.003, 12.A.006.

##### Etymology

From Velaux-La Bastide Neuve, Bouches-du-Rhône, the type locality.

##### Diagnosis

As for genus.

##### Description

see SI

#### III.1b: The associated vertebrates

Four of the brachyuran claws were discovered in close association with vertebrate remains. The most complete and larger specimen (MMS/VBN.00.004) is fossilized onto a vertebra that closely resembles a posterior cervical vertebra of a rhabdodontids Iguanodontia (Fig 1a–b). Rhabdodontids are represented at Velaux by the genus *Matheronodon* Godefroit et al., 2017^29^ and in other Late Cretaceous localities from southern France, by *Rhabdodon* Matheron, 1869^68^. MMS/VBN.09.69e is found in close sedimentary association with the plastral plate of a terrestrial turtle (*Solemys* de Lapparent de Broin & Murelaga, 1996^69^, Solemydidae) (Fig 1c–d). MMS/VBN.12.A.006 is preserved in a 50 cm large block that also contains a turtle plastral plate (*Polysternon* Portis, 1882^70^, Bothremydidae), a rhabdodontid tooth and centrum, as well as hybodontid shark teeth (Fig 1e–f). MMS/VBN.12.A.003, figured isolated (Fig. 3d), has been extracted from a comparable block, also containing a crocodylomorph skull, a rhabdodontid tooth, as well as a partial titanosaurid dorsal vertebra showing preserved ossified tendons.

### III.2. Freshwater environment

#### III.2a: Taxonomic/ecologies diversity in the sequence 2 of the channel

The freshwater palaeoenvironment of the channel in sequence 2 is strongly supported not only by sedimentological evidence^23^, but also by the most recently collected taxonomic assemblage itself. The fossil remains consist of 42.5% of strict terrestrial/aerian Avemetatarsalia indicative of an absolute continental faunal assemblage (Fig. 4). Aquatic and semi-aquatic taxa consist of families and genera, which previous depositional record is strongly anchored in freshwater environments. Among archosaurs, the hylaeochampisdae crocodylomorph *Allodaposuchus* Nopsca, 1928^27 71^) has so far been reported from fluvial inner/lacustrine-interpreted environments^69,72,73^ and once in a more coastal swampy area^74^. These are associated to a small amount of Globidonta, which, as members of Alligatoroidea, would have secondarily lost salt glands and therefore have been also restricted to freshwater settings^75^. Other highly abundant sauropsids in Velaux are chelonians, equivalently represented by Bothremydidae (*Polysternon*) and Solemydidae (*Solemys*, Fig. 4). If the former family is recognised as the most abundant and diverse European group of freshwater and coastal turtles in the uppermost Cretaceous^76^, *Polysternon* is only reported from estuarian to alluvial sediments and its sister-genus *Foxemys* Tong et al., 1998^77^ is exclusively known from freshwater localities^78,79^. The case of Solemydidae is even more compelling because their dermal skeleton (skull osteoderms) is highly supportive of a strict terrestrial life habit^80^ rather than any degree of amphibious lifestyle. The identified chondrichtiyan teeth correspond to a unique hybodontid genus: *Meristonoides* Case & Capetta, 2002^81^, which presence at Velaux has been briefly questioned (Cuny pers. comm. in^23^), although it is well accepted that hybodontid sharks are common in fluvial ecosystems in the Cretaceous^82^. The least abundant remains at Velaux belong to an aquatic sarcopterygian identified as *Axelrodichthys megadromos* Cavin et al., 2016^83^ (Cavin pers. comm.). The only known occurrence of this mawsoniid coelacanth is from another French Campanian lacustrine deposit^83^, confirming the unequivocal freshwater nature of the fauna from the sequence 2, which includes the brachyuran claws in the Velaux-La Bastide Neuve channel (Fig. 4).

**Figure 4.**
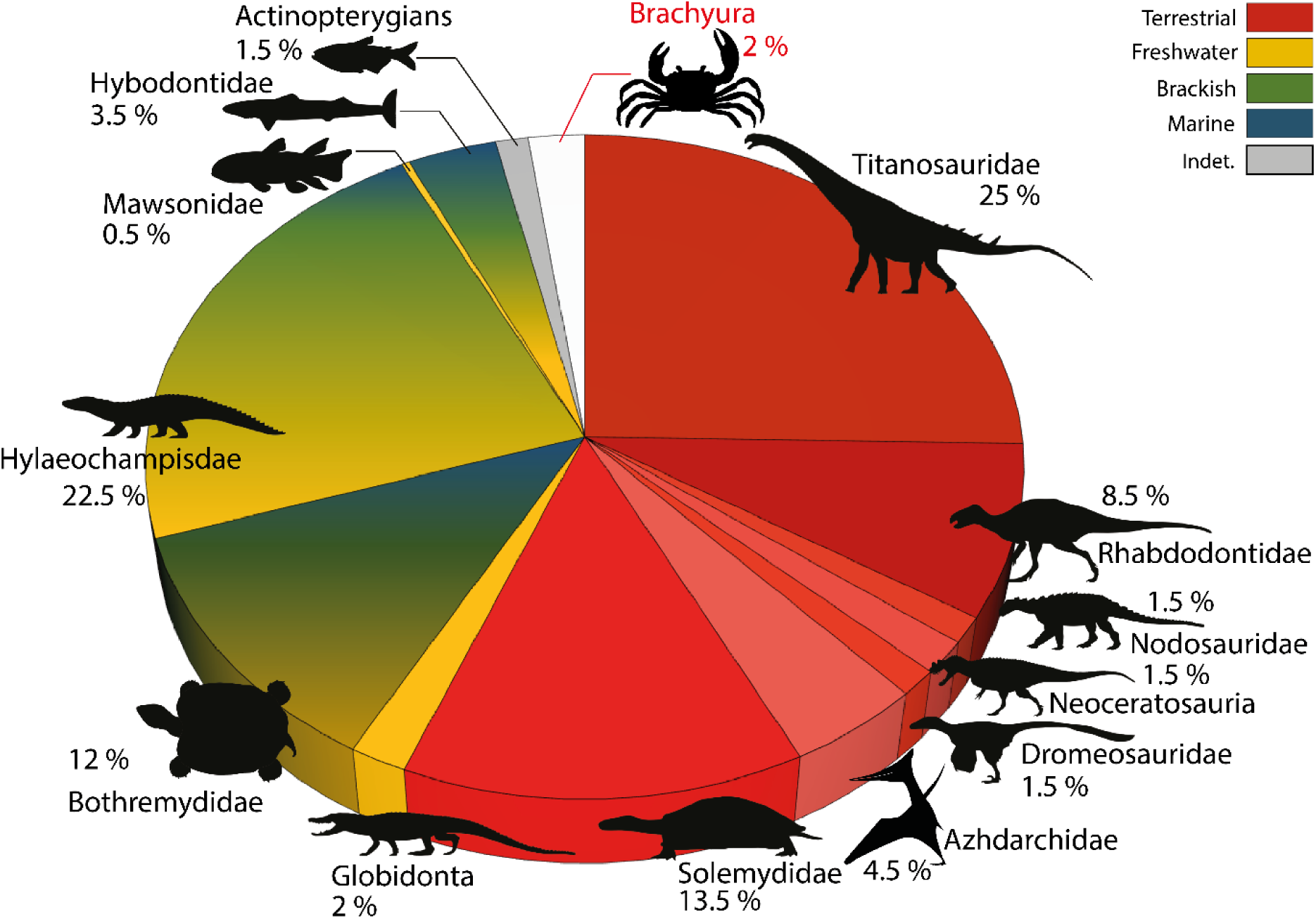
Distribution of the taxa remains, recovered from the sequence 2 and associated described depositional environments (see *III. 2a. Taxonomic/ecologies diversity in the sequence 2 of the channel*), based on Velaux-La Bastide Neuve excavation campaigns.

#### III.2b: Y/Ho ratios

As for other rare-earth-elements, the Y signature of limestones and carbonate concretions can be used as recorder of ancient seawater signatures^84–86^. Y and Ho concentrations are compared because these elements are chemically similar in charge and ionic radius, and suggested to evolve similarly in poorly terrigenous diagenetic environments^85^. The carbonates of MMS/VBN.09.69e-claw display a Y/Ho ratio of 33.06 (Y=8.61 ppm; Ho=0.26 ppm). The ratio in MMS/VBN.09.69e-sediment is a bit lower with 29.86 (Y=11.3 ppm; Ho=0.37 ppm). Marine waters are known to have a quite constant Y/Ho ratio around molar 90-110, decreasing with depth^87,88^. Consequently, neither the sandstones hosting the claws, nor the claw carbonate could have been deposited and/or formed in marine conditions. Apart from the marine realm, Y/Ho data characterizing formally typical estuarine or fluviatile environments are hitherto not much reported^89^. Nozaki et al.^89^ evidenced from the study of Japanese fluvial systems that Y and Ho concentrations were constantly decreasing with the salinity, with Ho removed from seawater twice as fast as Y owing to differences in surface complexation behavior. Unfortunately, Y and Ho absolute concentrations, which we would expect to interpret from the studied fossil/sediment material, depends on a biological and a taphonomic factor of integration, which cannot be estimated. Consequently, the actual salinity of the studied channel cannot be assessed from the chemistry but the observed Y/Ho ratio formally excludes a marine pattern and seems to distinguish strongly from it.

### III.3. Taphonomy of the assemblage

The crab chelae studied herein were recovered from the fluvial channel sediments from sequence 2, which correspond to lenticular conglomeratic sandstone (Fig. 5a-b). This association of elements belonging to diverse – aquatic and terrestrial – vertebrate taxa probably results from the transport of decayed carcasses originating from diverse environmental settings in a river channel. The associated bones and tooth elements are found disarticulated. The preservation of some complete bony elements, like crocodylomorph skulls, argues for their relatively short timing of decay and transport, consistent with the most common reports of the group in fluvial/ inner lacustrine type of environments. The Velaux taphonomy would indicate a local riverine system with a low-enough energy to allow the deposit of small millimetric elements like *Meristonoides* teeth^23^. An option could be that large elements (large appendicular bones, carapace portion and skulls) would have acted as obstacles for smaller ones in a more intermediate-energy flow configuration, resulting in a mixture of elements of different sizes and spatial origins^23^. In both cases, as in any continental water system, the deposit must have occurred up to downstream implying that all the remains in sequence 2 must have belonged either to original local fluvial living individuals (sedimentary context) or to upstream/even more terrestrial ones (floodplain and levees). Consequently, the presence of strictly freshwater lineages (Globidonta crocodylomorphs), would restrict the salinity inside this part of the channel to a minimum, implying that crabs must have been living either in terrestrial or freshwater aquatic habitats. The spatial distribution of the crabs within the conglomeratic sequence (sequence 2 on Fig. 5c-d) is heterogeneous: they are in most cases horizontally spaced by several dozens of centimeters. The absence of further connection of the brachyuran remains (e.g. with manus/carpus) or other body parts than dactyli and/or propodi is poorly informative on transport/exposure time experienced by claws given the admitted proclivity of decapod crustaceans’ chelae to preserve the best after years (see^90,91^ for brachyurans).

**Figure 5.**
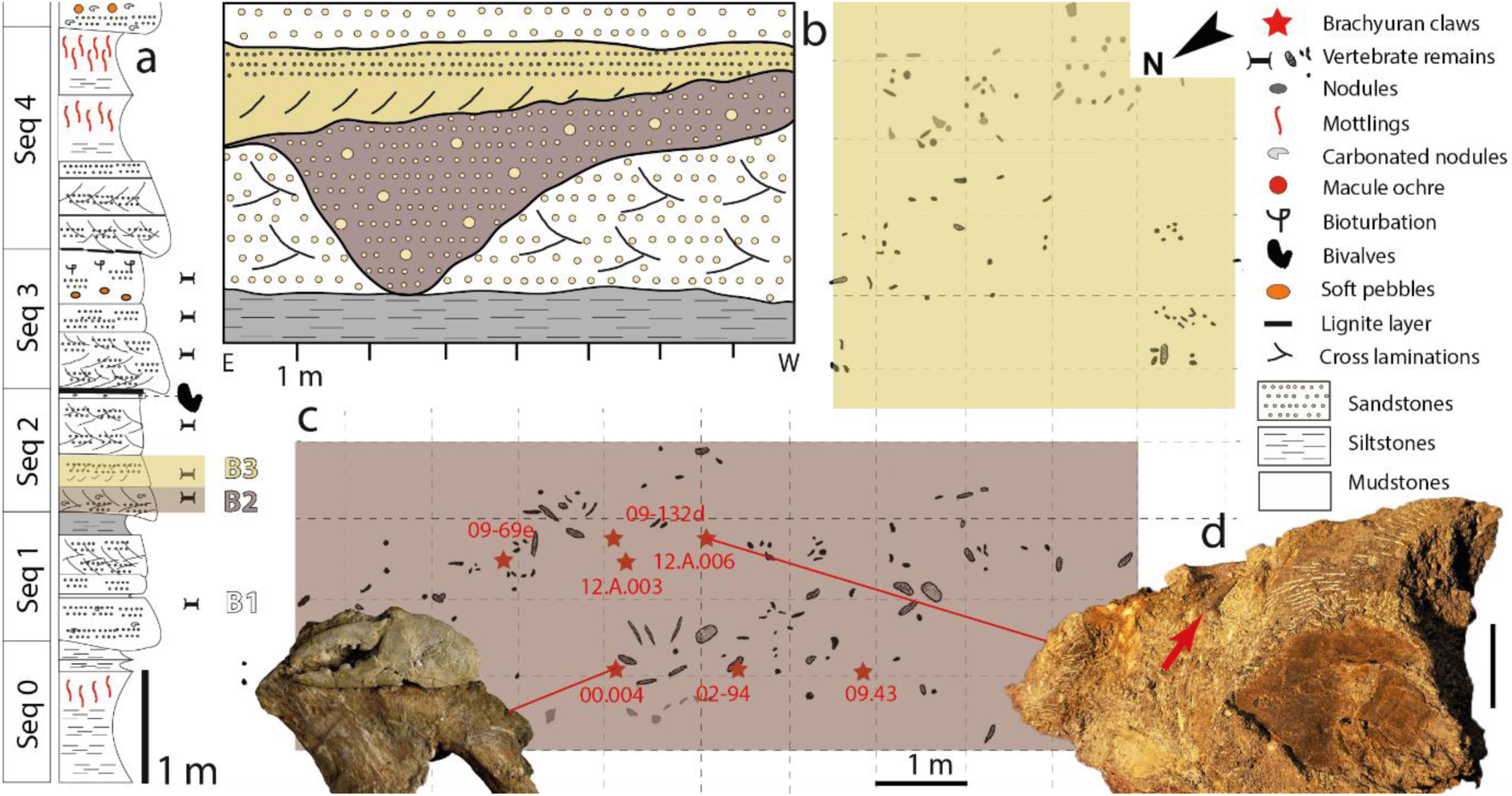
Distribution of the fossil assemblage in the deposit of Velaux-La Bastide Neuve, France. **a.** Lower portion of the Velaux log (Seq 0-4). **b.** Schematic spatial extension of the Velaux channel showing the distribution of bonebeds 2 (B2) and 3 (B3) in section. **c.** Top view of the distribution of claws/bones on the excavation site, the croping lower part of the sequence 2. **d.** Bloc assemblage of claws (small arrow) among vertebrate remains, scale bar = 10 cm (d). Modified from Cincotta et al. 2015 (a), field surveys from G. Garcia & X. Valentin (b, c), photographs. L. Cazes and X. Valentin (d).

## IV. Discussion

Beside their proposed systematics assignment, claw shape and important size cannot be used to inform on a specific diet or life habit of the crabs, nor on the actual interactions they could have maintained with reported members of their ecosystem. Interestingly, it is of note stressing that consumption behaviours of decapod crustaceans by megaherbivorous dinosaurs have been reported from the Campanian of two North American formations^92^. In that case, representatives of ankylosaurs, brachylophosaurs and neornithischians were point out to display scattered undetermined crustacean cuticles associated to fragments of likely rotted wood fragments in their in-situ coprolites. This led the authors to suggest these crustaceans were consumed when sheltered in dead logs without these cuticles could be identified as belonging to a specific order, although possibly corresponding fossil crab-like claws (not identified as Eubrachyura) were known from surrounding continental middle Campanian formations^92–94^.

### IV.I. Decapod crustaceans in freshwater habitats

Decapod crustaceans inhabit virtually all water-influenced habitats, including freshwater bodies, from streams and rivers to ponds and lakes, and even caves. In fact, representatives of a number of originally marine decapod clades have successfully invaded freshwater and/or terrestrial habitats. Among caridean shrimps, more than 650 species, making a full quarter of all described species, inhabit freshwater^95^, with representatives of *Merguia* Kemp, 1914^96^ being semi-terrestrial^97^. With approximately 650 species, virtually all crayfish are freshwater animals^98^. Many axiideans and gebiideans are able to tolerate pretty low salinity conditions. The callianassid *Lepidophthalmus* Holmes, 1904^99^ is even able to tolerate freshwater environments^100^ and *Lepidophthalmus turneranus* (White, 1861^101^) has been reported to migrate up rivers in West Africa^102^. Among anomurans, a rather speciose family Aeglidae Dana, 1852^103^ is strictly freshwater^104–106^. The majority of freshwater decapod crustaceans, however, consists of Brachyura with one fifth (>1 280 species) of all^107^.

Primary (so called *true*) freshwater crabs are those that have adopted freshwater, semi-terrestrial or terrestrial modes of life, and are able to complete their life cycle independently of the marine environment^1^. However, there are a number of brachyuran crabs able to live in freshwater habitats that include euryhaline species or secondary freshwater species from primarily marine brachyuran families^1^. These do not have direct development in their life cycle, which is typical for true freshwater crabs. Today there are five primarily freshwater families of brachyuran crabs^107^, i.e. Gecarcinucidae, Potamidae, Potamonautidae, Pseudothelphusidae, and Trichodactylidae, whereas there are also numerous secondary freshwater, semi-terrestrial and terrestrial species among Majoidea (Hymenosomatidae), Goneplacoidea (Goneplacidae), Grapsoidea (Gecarcinidae, Sesarmidae, Varunidae) and Ocypodoidea (Ocypodidae)^1^. Many grapsoids invade or even wholly inhabit freshwater habitats. Some varunids, including representatives of *Eriocheir* De Haan, 1835^108^ and *Varuna* H. Milne Edwards, 1830^109^, not only enter estuaries, but are also found further up in rivers^1^. Sesarmids (*Sesarmoides*, *Labuanium*, *Karstarma*) can be completely adapted to freshwater, the latter being semi-terrestrial^107,110–112^, whereas *Geosesarma* De Man, 1892^113^ is found in terrestrial habitats^110,111^.

As for the fossils from Velaux and surroundings newly described herein, it cannot be decided unequivocally whether representatives of *Dinocarcinus* n. gen. were able to complete their life cycle in the freshwater habitat or/and had direct development. The sheer size and number of claw fragments, however, may prove that crabs from Late Cretaceous of Velaux were not only an occasional element of the respective environment, but rather a natural part of the assemblage suggesting that they were fully adapted to freshwater environment.

### IV.2. Freshwater decapods in the fossil record

Fossil freshwater decapods are exceedingly rare in comparison to their marine relatives. Fossils crayfishes are represented only by a handful of occurrences reported so far (^110,111,114–118^ and references therein), the oldest coming from the Triassic of Utah^115^. Fossils of freshwater caridean shrimps are similarly rare^114,116,119–122^, the oldest being reported from the Early Cretaceous of Spain^116^ and China^122^. Interestingly, the only fossil representative of nowadays strictly freshwater anomurans, the family Aeglidae, comes from marine strata^123^. Fossil freshwater brachyurans are limited to a number of occurrences of isolated claws^9,15–18^ and several fossils exhibiting preserved carapace^7–13,124^. The oldest occurrences of undisputed primary freshwater crabs are from the middle Eocene (late Lutetian/early Bartonian) of the Amazon Basin, recently reported by Klaus et al.^7^ who described isolated claw elements of Trichodactylidae, and from the middle Eocene (Bartonian) of Italy (*Alontecarcinus buratoi* De Angeli & Caporiondo, 2019^124^) being the oldest representative of Potamidae. In this respect, *Dinocarcinus velauciensis* n. gen. n. sp. reported herein is the oldest occurrence of freshwater brachyuran crabs, exceeding previous reports by approximately 40 million years. For now, it is unclear whether *Dinocarcinus* belonged to primary or secondary freshwater crabs. It is, however, of note that recent advances in resolving the phylogeny of primary freshwater crabs suggest their early divergence in brachyuran evolution^125^.

### IV.3. Multiple invasions into freshwater habitats

From the discussion above it is clear that several lineages of decapod crustaceans independently invaded freshwater habitats, including dendrobranchiatans^126^, carideans, axiideans, astacideans, anomurans and brachyurans^6,19,105,127^. The enigmatic *Tealliocaris* Peach, 1908^128^, considered by some authors as a decapod crustaceans^129^ (but see also^130^), might represent yet another freshwater lineage. And among carideans, at least palaeomonoid, atyoid and alpheoid shrimps independently invaded freshwater environments^95^. Moreover, presumed multiple invasions of freshwater habitats by some *Macrobrachium* Bate, 1868^131^ shrimps were also suggested^132^. The sparse fossil record of freshwater shrimps does not allow relevant time estimation of colonization of freshwater habitats; however, fully freshwater shrimps are known from the Early Cretaceous (Barremian) onward^116,122^. Crayfish represent a monophyletic group^133^ with the oldest fossil representatives known from the Late Triassic of Utah^115^. As for brachyuran freshwater crabs, there are two independent lineages. The Old World primary freshwater crabs are monophyletic^6,19,107,134^, whereas Neotropical Trichodactylidae have a separate phylogenetic origin and appear to be closely related to marine Portunidae^19,135^. Based on the morphology of its chelae, *Dinocarcinus velauciensis* n. gen. n. sp. cannot be referred to any of the extant primary or secondary freshwater families mentioned above suggesting that it represents yet another independent “attempt” to colonize freshwater environment besides the two primary freshwater crab clades recognized today, i.e., Potamoidea and Trichodactyloidea^19,107^. Interestingly, the oldest fossil representatives of both clades come from the middle Eocene^7,124^. The geographic distribution of modern primary freshwater crabs speaks for independent invasions of the limnic habitat rather than for a Gondwanan vicariance^6,136^, contrasting with the diversification of crayfishes: the fossil record and modern distribution of the latter clade can be explained by the breakup of Pangaea and disassembly of Gondwana and Laurasia^127^. Based on molecular clock estimates, Daniels et al.^136^ suggested that the radiation of Afrotropical freshwater crab taxa occurred during the Early Cretaceous, whereas the age of the African Potamonautidae clade was given with 75-73 Ma (Campanian). From the fossil record of the modern freshwater families alone, such timing cannot be apprehended; however, the discovery of fossil crabs from Velaux-La Bastide Neuve illustrates that brachyuran crabs attempted to colonize freshwater habitats in the Old World at least from the Campanian onwards.

One of the key processes driving freshwater crab diversification is likely allopatric speciation resulting from geographic isolation, often coupled with habitat heterogeneity and numerous ecological niches and microhabitats resulting from the complicated topography and hydrology of freshwater environments^1^. During the Campanian, *Dinocarcinus velauciensis* inhabited Europe, which was at its time an archipelago rather than a proper landmass^32,137^. Based on the material from Velaux and Rognac described herein, we suggest that the freshwater habitats of islands in the Tethyan epicontinental sea were colonized by marine portunoids during the Late Cretaceous. Nowadays, most secondary freshwater brachyurans have a marine larval development and would reach inland habitats more likely as adults. This might also have been the case for *Dinocarcinus velauciensis*.

## V. Conclusions

*Dinocarcinus velauciensis* n. gen. n. sp. from the late Campanian of Southern France, belongs to Portunoidea sensu lato, a group of “true crab” that are nowadays intimately linked to marine systems. The sedimentological context, faunal assemblage and taphonomy of these fossils, as well as the Y/Ho ratio of their carbonates indicate an ancient freshwater or terrestrial ecology. This make them the oldest freshwater/terrestrial brachyurans ever reported, extending the existence of freshwater crabs by 40 Ma. In this Campanian ecosystem, “true” crabs were intimately associated to terrestrial vertebrates, including non-avian dinosaurs. Although they were likely well adapted to this environment, it cannot be decided whether *Dinocarcinus* was able to complete its life cycle in the freshwater habitat or/and had direct development. Its occurrence in the Late Cretaceous of Velaux-La Bastide Neuve, is an evidence for the independent colonizations of freshwater environments by multiple Brachyura clades over time, beside that of modern primary freshwater crabs (Potamoidea, Trichodactyloidea). It also supports the molecular clock estimation of an Early Cretaceous start for the radiation of Afrotropical freshwater crab taxa (just appearing in the Late Cretaceous), with the evidence of brachyuran crabs colonizing freshwater habitats as early as the Campanian.

## VI. Methods

The elementary composition of a brachyuran claw and of its surrounding matrix were investigated for their Y/Ho ratios. One gram of each was sampled on MMS.VBN.09.69e (claw/sediment). For the claw material, the basis of the propodus embedded in the matrix was mechanically sampled to preserve the connection between the claw and the turtle plate. Samples were microgrinded and analysed for their composition in minor elements (in µg/g) normalized to PAAS, using ICMPS at the *Service d’Analyses des Roches et des Minéraux* of the CRPG, Vandoeuvre-lès-Nancy, France.

## Data Availability

All data needed to evaluate the conclusions in the paper are present in the paper and supplementary information.

## VII. Acknowledgments

Authors want to acknowledge Clément Jauvion for discussions on elementary ratio in marine/non marine sediments. XV and GG thank Y. Dutour and E. Turini for photographs of the specimen from Rognac. GG and XV gratefully acknowledge the management, logistical, and communication assistance from the Velaux Municipality (J.-P. Maggi and L. Melhi) with its heritage, culture, and technical services (M. Calvier and S. Chauvet), the environment department from CD 13 (M. Bourrelly, T. Tortosa, G. Michel, N. Mouly, and S. Amico), the Service Départemental d’Incendie et de Secours (SDIS) 13, and numerous volunteers during the field campaigns in 2009 and 2012. This work was supported in part by the French Ministry of Culture and Communication (research grant VR1013 to the Palaios Association) and the Bouches-du-Rhône department (CD 13) (proposals MAPADGAC23112010-1 and MAPADGAC16012014-1-AAPC).

## IX. Authors contributions

NR, BVB and MH performed the claw study, analyses and manuscript draft. AC, XV and GG provided the geological and taphonomic context of claws location. GG, PG and XV collected the fossils and provided data on faunal assemblage for Velaux deposit. SC supported material imaging and provided taxonomic review and comments. X.V. and G.G. supervised the project.

## X. Competing interests

The author(s) declare no competing interests.

