## Supplementary Information for "The oldest freshwater crabs: claws on dinosaur bones"

Portunoidea sensu lato (see *Prelimiary remarks*)

*Dinocarcinus* n. gen. Van Bakel, Hyžný Valentin & Robin

**Etymology:** Denoting the actual association with dinosaur (ornithopodan) remains.

**Type species:** *Dinocarcinus velauciensis* n. gen., n. sp.

**Diagnosis:** Chelae large and massive. Fingers gaping, arched, with strong teeth, proximal tooth molariform. Fixed finger dorsal surface with single "pitted groove", palm surface smooth, articulation with dactylus oblique, prominent.

*Dinocarcinus velauciensis* Van Bakel, Hyžný, Valentin & Robin n. sp.
Figs. 1, 2, 3

**Type material:** Holotype: MMS.VBN.00.004; Paratype 1: MMS.VBN.02.94; Paratype 2: MMS.VBN.09.132d; Paratype 3: MMS.VBN. 12.A.003; Paratype 4: MMS.VBN.12.A.006.

**Etymology:** From Velaux-La Bastide Neuve, Bouches-du-Rhône, the type locality.

**Diagnosis:** As for genus.

**Description:** Only claws known; claw very large (approximately 85 mm for holotype MMS.VBN.00.004), massive, outer surface flat. Palm subrectangular, slightly longer than high, slightly longer than fixed finger. Fixed and movable fingers inwards curved, clearly gaping. Lower propodus margin curved, weakly convex. Upper (cutting) margin of fixed finger straight. In total, 4 strong teeth on fixed finger. Proximal tooth massive (T1/t1), molariform in both fingers. Surface of molariform proximal tooth bulbous on fixed finger, flat on dactylus. Fixed finger dorsal surface with a single "pitted groove". Articulation dactylus-propodus prominent, oblique. Both finger tips sharp, pointed. Dactylus upper and lower margin curved in dorsal view; dactylus cross-section flatter than propodus cross-section. Dactylus more strongly curved in dorsal view than fixed finger. Dactylus margin at approximately 100 degrees at upper (cutting) margin of fixed finger. Fingers strongly calcified, cuticle surface (as preserved) covered with microscopic dense, flattened granules.
